# Selective Cell Propagation via Micropatterning of Thermal-activated Hydrogel

**DOI:** 10.1101/2020.04.01.019208

**Authors:** Jeffrey C.Y. Chiu, Joyce A. Teodoro, Jeong Hyun Lee, Kerryn Matthews, Simon P. Duffy, Hongshen Ma

## Abstract

The ability to selectively propagate specific cells is fundamentally important to the development of clonal cell populations. Current methods rely on techniques such as limiting dilution, colony picking, and flow cytometry to transfer single cells into single wells, resulting in workflows that are low-throughput, slowed by propagation kinetics, and susceptible to contamination. Here, we developed a method, called selective laser gelation (SLG), to micropattern hydrogels in cell culture media in order to encapsulate specific cells to selectively arrest their growth. This process relies on the inverse gelation of methylcellulose, which forms a hydrogel when heated rather than cooled. Local heating using an infrared laser enables hydrogel micropatterning, while phase transition hysteresis retains the hydrogel after laser excitation. As a demonstration, we used this approach to selectively propagate transgenic CHO cells with increased antibody productivity. More generally, hydrogel micropatterning provides a simple and non-contact method to selective propagation of cells based on features identified by imaging.

**One Sentence Summary:** Inverse gelation of methylcellulose enables hydrogel micropatterning to selectively propagate cells identified by microscopy.

## Introduction

Biological systems are inherently heterogeneous, and this heterogeneity has driven a longstanding need to isolate and propagate individual cells to develop clonal populations. The development of clonal populations has contributed to tremendous advances, such as identification of disease-causing agents^1^, screening for potential therapeutics^2,3^, delineation of the roles of cell subpopulations within tissues^4^, as well as development of cell lines for biologics production^5^. The generation of monoclonal cell populations remains a challenging and laborious pursuit. The key challenges include long lead-times needed to grow single cells to significant number before they can be screened for desirable phenotypes, technical challenges associated with isolating and transferring single viable cells, as well as the potential that contact with the extraction instrument may contaminate the isolate. For these reasons, the development of clonal populations remains a laborious process with a high rate of attrition.

One common context for single cell isolation and cloning is the generation of high yield monoclonal antibody production systems. Genetic engineering has yielded highly efficient recombinant producer cells^6^, however the introduction of immunoglobulin transgene can contribute to cell-to-cell variability in antibody secretion, owing to natural variability in host cell metabolism, as well as imprecise positioning of the transgene and epigenetic silencing of the antibody-producing loci for recombinant antibodies^7^. This variability in host cell productivity can be magnified over time during scale-up production where fast-growing cells are propagated at the expense of high-producing cells, which often grow at a slower rate, thereby limiting the ultimate productivity of the bioreactor^8^. Consequently, additional screening is required to develop host cell lines that maximize productivity, while at the same time minimize heterogeneity prior to scale-up production.

Current strategies for developing high-producing cell lines for antibody production include limiting dilution, colony picking, fluorescence activated cell sorting (FACS), gel microdroplet sorting, and laser-enabled analysis and processing (LEAP). Limiting dilution and colony picking are both laborious and time-consuming methods that require propagating single cells into colonies that are sufficiently large to have their production rate measured using ELISA-based quantification^9–12^. These methods screen producing cells at the level of colonies, which may be already dominated by fast-growing, but low-producing cells. Alternatively, screening at the single cell level based on cell surface expression could be performed using FACS. However, this approach is unreliable since cell surface expression often does not correlate with antibody secretion rate^5^. Gel microdroplet sorting overcomes this issue by encapsulating individual antibody producing cells in a microdroplet of agarose. The droplet contains reagents for fluorescence labeling of secreted antibodies, enabling FACS sorting of droplets containing high-producing cells^13,14^. Challenges associated with this technique include the need to optimize the single cell encapsulation process for each cell line, as well as the concern that liberating cells from the gel microdroplet via agarose digestion could adversely affect host cell viability^13,15^. Finally, LEAP is a proprietary process that captures host cells and secreted antibodies on a functionalized surface, in order to selectively lyse low-producing cells using a focused laser, to enable the propagation of high-producing cells^16^. While this technology has demonstrated potential to selectively propagate high-producing cells, the adoption of this technology has been hindered by the cost and complexity of the system, as well as the need for custom reagents and disposables^12^. Therefore, despite the variety of available screening methods, there remains a significant need for a process to selectively expand high-producing cells prior to scale-up production.

Recently a number of approaches have been developed to separate cells based on imaging by selectively immobilizing cells on the surface of an imaging substrate using photo-polymerizable hydrogels or photo-activated surface chemistry. In approaches involving photo-polymerizable hydrogels, cells are first adhered to a surface by culturing or capturing in nanowells. The undesired cells are then selectively entrapped on the substrate by encapsulation using photo-polymerizable hydrogel, while target cells can then be extracted by washing^17,18^. A similar strategy has been deployed by culturing cells or specifically capturing cells on a photo-degradable hydrogel substrate, where target cells can be removed by selective dissolving the hydrogel underneath target cells^19–22^. In approaches involving photo-activated surface chemistry, cells are cultured on a substrate. Unwanted cells are adhered to the substrate by photo-crosslinking, while target cells are removed by enzymatic digestion^23,24^. A major challenge for using these approaches to isolate clonal cells for antibody production is that they do not integrate the ability to measure secretion from each cell in order to identify producing cells and non-producing cells. Additionally, these methods often require exposure to chemicals agents (e.g. hydrogel pre-polymers) that have some degree of toxicity to cells^17^. Although, short-term exposure to these agents can usually be tolerated by immortalized cells, growing cells in the presence of these chemical is typically undesirable. Therefore, target cells must be extracted and transferred at the stage of one or a few cells, which is prone to loss and contamination.

Here, we describe a new technology for developing clonal populations of cells by selectively arresting the growth of unwanted cells in a process called Selective Laser Gelation (SLG). SLG relies on the inverse gelation of methylcellulose in order to micropattern hydrogels to encapsulate specific cells to arrest their growth. The hydrogel is retained after removing the laser due to gelation hysteresis, enabling outgrowth of desired cells. Importantly, this process provides an image-based selection method to specifically propagate target cells, but does not require extracting or transferring single cells, which dramatically improves throughput and reduces the potential for contamination and loss of viability. We demonstrate that SLG is capable of enabling selective growth of high-yield antibody producing CHO cells, but this capability can also be applied to a wide range of applications where the generation of clonal cell populations is needed.

## Results

### Material Properties of Methylcellulose

Methylcellulose is a common reagent used to provide a semi-solid matrix to immobilize producer cells and localize antibody secretion prior to imaging and colony picking^11,25^. It is also a well-known non-toxic additive used for culturing stem cells, typically administered at 1% working concentration^26^. Secreted antibodies are detected in methylcellulose solution using fluorescence-labeled anti-IgG^27^, which form aggregates with secreted antibodies resulting in a fluorescent halo around each producer cell^28^. The size and brightness of the halo can provide a measure of the antibody secretion rate of each producer cell.

Cell culture media supplemented with methylcellulose has the unusual property of forming a hydrogel when heated rather than when cooled^29,30^. This sol-gel transition is also hysteretic, where the solution-to-hydrogel transition temperature is greater than the hydrogel-to-solution transition temperature^31,32^. The sol-gel transition temperatures and hysteresis band depend on methylcellulose concentration. Specifically, we determined that a methylcellulose concentration range of 1 to 2% results in a material with a gelation temperature greater than 37°C and a dissolving temperature less than 37°C (**Figure 1**). To demonstrate this property, we heated a vial of 2% methylcellulose solution from 20°C to 60°C, which caused the solution to transition to gel phase (**Figure 1A**). As the vial was cooled to 37°C, the methylcellulose solution remained in the gel phase (**Figure 1B**). Further cooling to 20°C resulted in the dissolution of the gel and restoration of the solution phase (**Figure 1C**). Increasing the vial temperature back to 37°C resulted in retention of the solution phase (**Figure 1D**). The combination of inverse sol-gel transition and gelation hysteresis (**Figure 1E**) enables (1) patterned gel formation using localized laser heating, as well as (2) the gelled regions to remain after removing the heat source.

**Figure 1.**
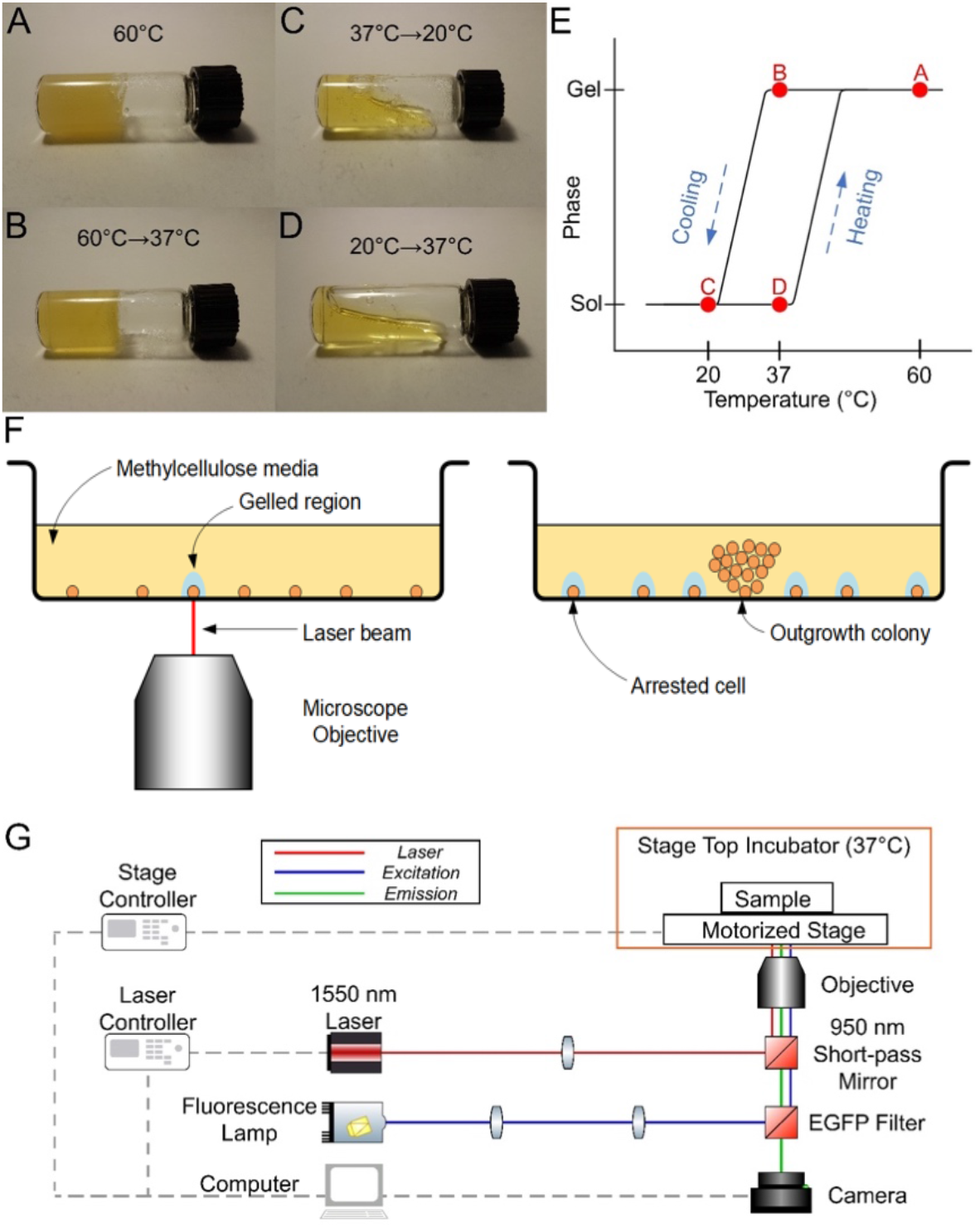
Principles of Selective Laser Gelation. **A-E:** Material properties of methylcellulose enabling selective laser gelation. Methylcellulose solutions form a hydrogel when heated and melts to solution when cooled. This phase change is hysteretic around 37°C for a solution containing 1% methylcellulose. **F:** The principle of Selective Laser Gelation (SLG) where localized laser heating selectively encapsulates target cells in hydrogel to arrest their growth, while non-gelled cells are allowed to propagate. **G:** The experimental apparatus used to test the SLG process.

### Selective Laser Gelation

Based on the unique inverse sol-gel transition property of methylcellulose, we developed a process to selectively control cell growth by using a focused infrared laser beam to micropattern hydrogels in methylcellulose media in order to selectively encapsulate single cells or cell colonies to restrict their growth (**Figure 1F**). The experimental apparatus consists of an inverted microscope outfitted with a camera, motorized stage, stage-top incubator, and two epi-fluorescence imaging ports. The stage-top incubator is used to maintain the methylcellulose media temperature at 37°C, which is crucial in selective hydrogel formation via focused laser application. One of the epi-fluorescence ports is used for standard fluorescence imaging, while the second epi-fluorescence port is used to introduce a 1550 nm laser into the optical path via a 950 nm short-pass mirror (**Figure 1G**). The diameter of the resulting gelation region is a function of the amount of energy absorbed by the laser, as well as the localized distribution of this heat through thermal conduction. We selected the 1550 nm laser to locally heat methylcellulose because of its compatibility with microscope optics and its high energy coupling to water. The operation of the laser, microscope stage, and camera were controlled by computer software, which targets the laser based on the location of cells determined using the microscope camera.

The custom computer software controlling the camera, motorized stage and laser was implemented in C# with the user interface created using Windows Presentation Foundation (**Figure S2-3**). The Main Control tab of the user interface establishes connection with the stage, laser controller and camera, and allows for laser and camera parameter fine-tuning (**Figure S3A**). The Stage tab controls the scanning and image stitching functions, which are necessary when imaging wells that do not fit within the field of view of the camera and microscope objective (**Figure S3B**). It also displays the current stage position and presents an option to manually move the stage to a desired position. Diagnostic tools for troubleshooting serial communication with the motorized stage and laser are contained in the Serial tab (**Figure S3C**). Once a target well is scanned and imaged, the resulting stitched brightfield and fluorescence images are processed through the Image control panel (**Figure S3E**), as described in the pipeline illustrated in **Figure S5**. The image processing pipeline produces a list of cell colony locations for laser targeting expressed in terms of pixel values, which are converted to stage coordinates in the Position panel (**Figure S3D**).

The SLG screening process begins by suspending cells in 1% methylcellulose, which is achieved by diluting 2.5% methylcellulose with culture media. CloneDetect is also added to the solution, which is a reagent containing fluorescence-labeled anti-IgG. The CloneDetect reagent labels and aggregates secreted antibodies to enable them to be detected by fluorescence microscopy. The cell suspension is seeded into a 96-well flat-bottom imaging well plate and cultured for 1-7 days in an incubator. The well plate is then imaged using the inverted microscope to identify cells and measure their antibody secretion. After identifying target cells, the SLG process is used to encapsulate all non-target cells in order to selectively arrest their growth.

### Localized Gelation and Selective Arrest of Cell Growth

We initially tested the ability of the SLG process to form localized hydrogels around individual cells to arrest their proliferation. We seeded B13-24 CHO cells (IgG4-secreting cell line) in 1% methylcellulose in a 96-well flat-bottom microtiter plate. We imaged the cell suspension using a 20X objective, and then fired the 1550 nm laser at different spots in the well using 300 ms laser pulses in a range of powers. Gelled spots begin to appear when the laser power exceeded 500 mW (2500 mA). These spots were not visible using standard imaging, but could be seen as a faint circle under phase contrast imaging (**Figure 2A**). The diameter of the gelled spots increased with increasing laser power as expected (**Figure S1A**). The minimum gel diameter that could be achieved was 103 ± 9.02 μm (**Figure 2A and S1B**). We followed the growth of cells embedded in the gelled regions (**Figure 2B and 2C**). After 24 hours, cells in the liquid region of the semi-solid media continued to divide, while cells in the gelled region did not divide (green and yellow circles in **Figure 2B and 2C**, respectively).

**Figure 2.**
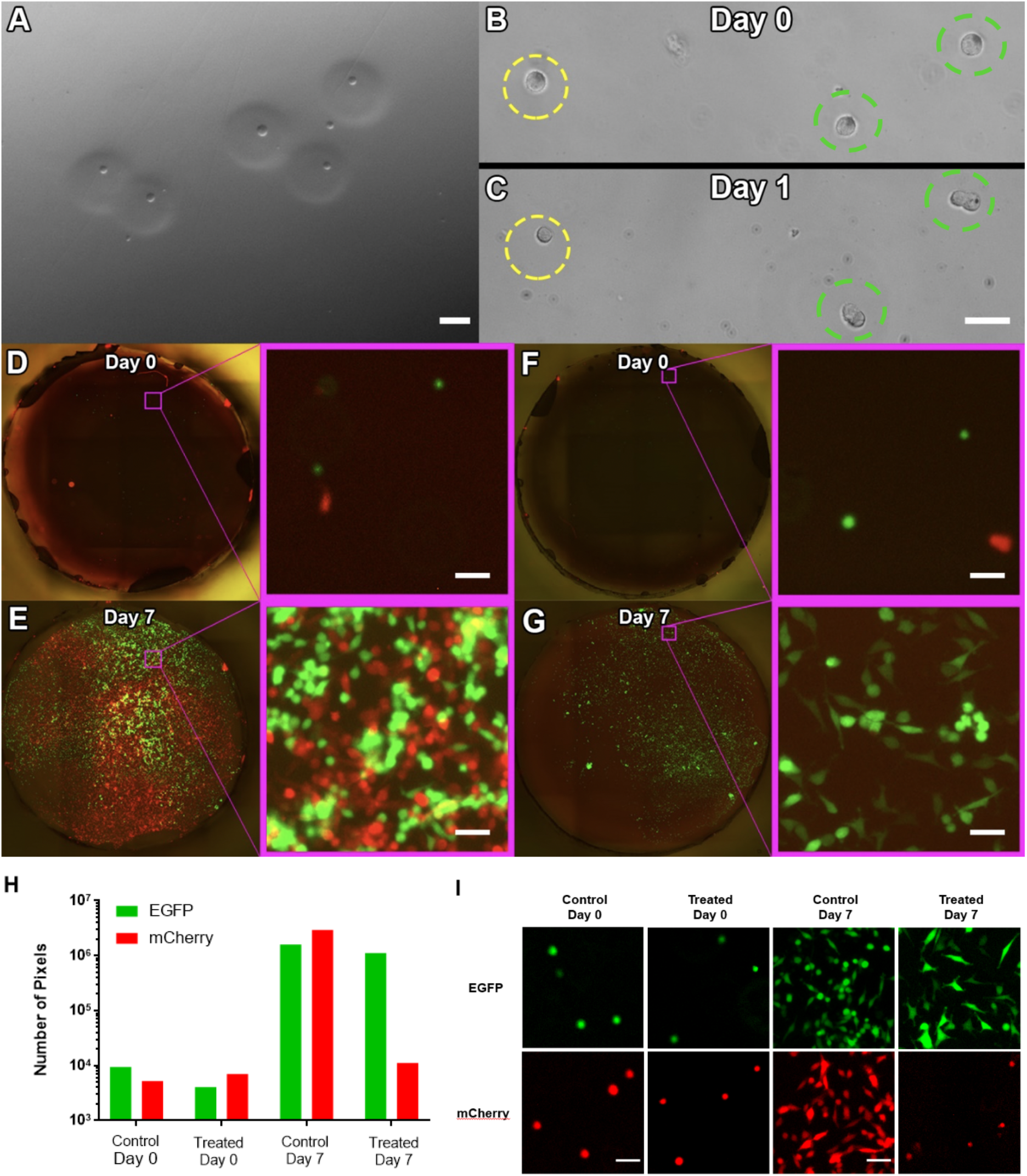
Validation of the SLG process. **A:** After SLG, gelled spots in methylcellulose are visible as faint circles using phase contrast microscopy. **B-C:** Growth of SLG-encapsulated cells (yellow broken circle) is restricted, while neighboring cells can proliferate (green broken circle). **D-E:** Co-culture of UM-UC3-mCherry (red) and UM-UC13-EGFP (green) produced a mixed population after 7 days. **F-G:** Using SLG to selectively eliminate UM-UC3-mCherry cells allowed only UM-UC13-EGFP cells to propagate after 7 days. **H:** Pixel count for mCherry and EGFP channels for control wells (D-E) and SLG wells (F-G) on days 0 and 7. **I:** UM-UC3-mCherry and UM-UC13-EGFP cell morphology and density on days 0 and 7 in the control and SLG wells. (Scale bars = 50 μm).

We then tested the SLG process at the scale of a well plate to selectively arrest the growth of one cell phenotype in a mixture. We generated a 1:1 mixture of transgenic UM-UC3 and UM-UC13 cells that expressed mCherry and EGFP respectively. We plated these cells in 1% methylcellulose in 96-well flat-bottom well plates (**Figure 2D and 2F**). In the control well without SLG, both cell lines propagated normally, such that large numbers of mCherry and EGFP cells could be observed after 7 days (**Figure 2E**). When SLG is used to selectively gel the mCherry cells, only the EGFP cells were able to propagate, while mCherry cells did not propagate after 7 days (**Figure 2F and 2G**). To quantitatively measure the propagation of UM-UC3-mCherry and UM-UC13-EGFP cells in each well, we analyzed the well images in **Figures 2D-G** by counting the number of red and green pixels (**Figure 2H**). Upon initial seeding, each well had 4 × 10^3^ to 9 × 10^3^ pixels in both mCherry and EGFP channels. After 7 days of propagation, the control well had mCherry and EGFP pixel counts increase to just under 3 × 10^6^ and 1.5 × 10^6^, respectively. The SLG well had only the EGFP pixel count increase to 1 × 10^6^, while the mCherry pixel count saw a modest increase to 1.1 × 10^4^ pixels. This slight increase is likely due to cell lysis, which resulted in spreading of the fluorescent cell debris. Morphologically, upon initial seeding, the UM-UC3-mCherry and UM-UC13-EGFP cells exhibited rounded morphology with smooth boundaries (**Figure 2I**). After 7 days, the propagated cells in the control well adhered to the well plate and showed spindle-shaped morphology. In the SLG-treated well, only the UM-UC13-EGFP cells attached and showed spindle-shaped morphology, while the UM-UC3-mCherry cells became fragmented with rough edges due to cell lysis. Together, these results demonstrate the ability to selectively arrest the growth of specific cells using the SLG process. Arrest of cell growth is a combination of cell lysis induced by local heating, as well as encapsulation in methylcellulose hydrogel, which deprives cells of nutrients for survival.

### Image Analysis to Measure Antibody Secretion

In semi-solid cloning, antibody secretion from producing cells is detected using a fluorescent secondary antibody, which forms aggregates with secreted antibodies^33^. The semi-solid matrix prevents the diffusive movement of these aggregates, which appear as fluorescent spots surrounding each cell or cell colony. To quantitatively evaluate antibody secretion, we developed custom software to measure and order the colonies based on the amount of secretion per cell. Specifically, this software captured a panel of both brightfield (BF) and green fluorescence images spanning the entire imaging plate well. The images from each channel were then digitally stitched into a single image. An image processing pipeline was implemented in software using EmguCV, a .NET wrapper for the Open Source Computer Vision Library (OpenCV)^34^. Using the BF image, cell colonies are detected using the Canny edge detector^35^ with the upper and lower thresholds automatically determined using Otsu’s method^36^. A contour detection algorithm^37^ is used to detect the centroid location (X, Y) and create a bounding box around the cell colony. The fluorescence image is run through an adaptive threshold to detect secreted antibody. Each centroid and bounding box area (*A_colony_*) detected from the BF image is used to count the number of pixels of secreted antibody (*N_ab_*) detected after the adaptive threshold from the fluorescence image. The colonies are then ranked using the ratio *N_ab_*/*A_colony_* (**Figure S4** and **Table S2**). The top ranked colonies are preserved and the remaining (X, Y) locations of centroids of the remaining colonies were used to move the motorized stage to center the laser on each undesired colony.

### Using SLG to Improve Antibody Secretion Rate

We tested the potential to use the SLG process to selectively propagate high-yield antibody-secreting cells using the B13-24 CHO cell line. B13-24 cells secretes anti-CD18 IgG4 at an average rate of 25 pg/cell/day^38^. Our process begins by suspending B13-24 cells in IMDM media with 1% methylcellulose and fluorescence-conjugated secondary antibody. The suspension was plated into the interior wells of a 96-well polystyrene well plate (**Figure 3A-B**). Following 1-2 days of limited culture, we imaged the well and used our software to determine the size, locations, and antibody production rate of each cell or colony using its surrounding fluorescent antibody aggregates (**Figure 3C-D**; **Figure 4A-E**; **Figure S3-5**). The cell colonies were then ranked based on the amount of antibody produced relative to the size of the colony (**Table S2**). In each well, we selectively retained either the three highest-producing colonies, or three lowest-producing colonies in separate wells. The remaining colonies were gelled by SLG to arrest their growth. This work was done automatically using our software to target the laser by controlling the motorized microscope stage (**Figure 3E**).

**Figure 3.**
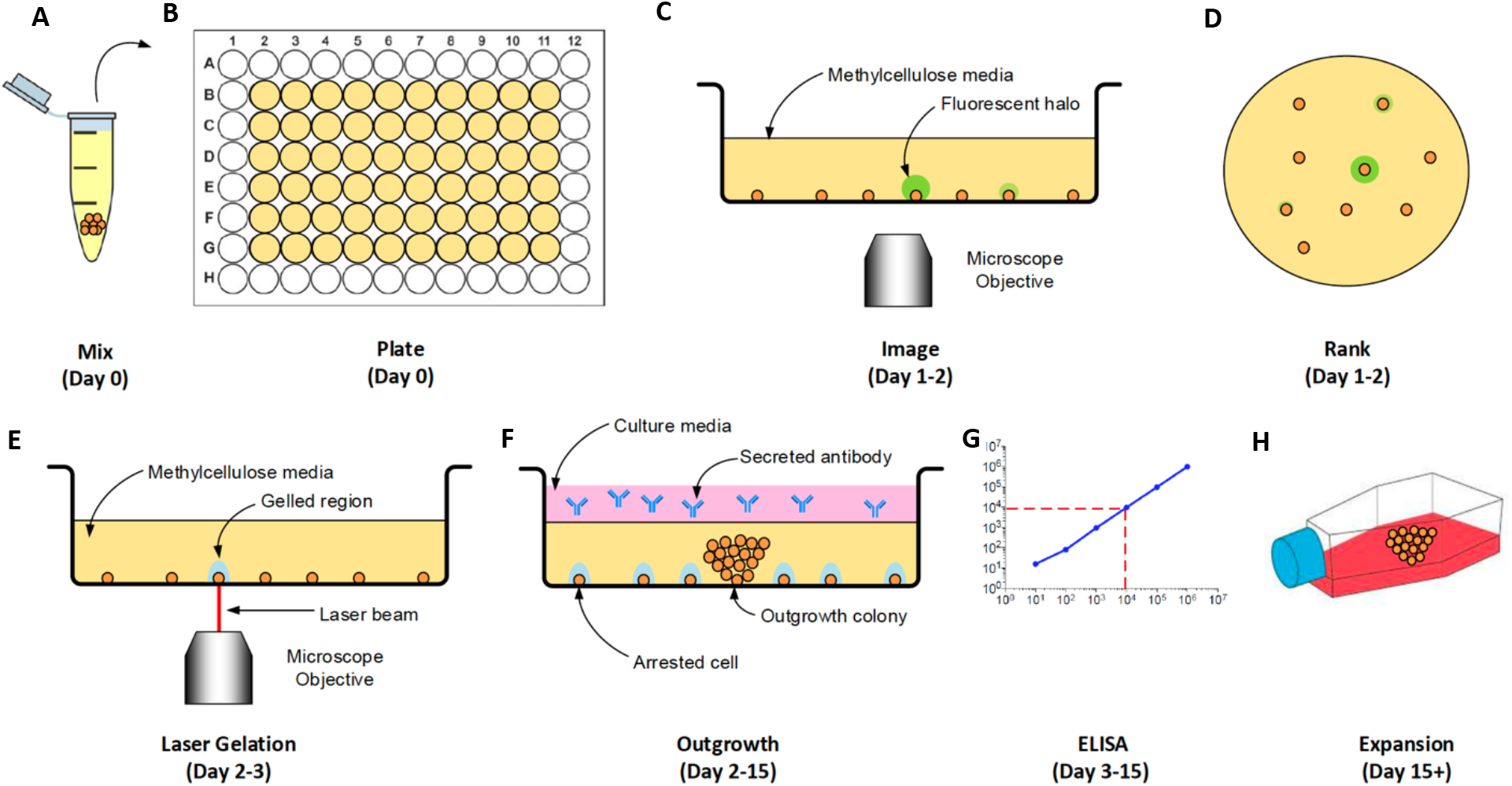
Process to select for high-producing CHO cell clones using SLG. **A:** CHO cells are mixed into a media solution containing 1% methylcellulose and 1% CloneDetect to detect secreted IgG4. **B:** The suspension is plated into a 96-well plate. **C:** Each well is scanned in bright field and fluorescence. **D:** Colonies are ranked based on antibody secretion per cell. **E:** The top-producing cells were retained, while SLG encapsulates the remaining cells in hydrogel to arrest their growth. **F:** Liquid media is added and the remaining cells are allowed to propagate. **G:** Antibody secretion rate can be measured via ELISA by sampling the supernatant. **H:** The culture is expanded using large containers.

**Figure 4.**
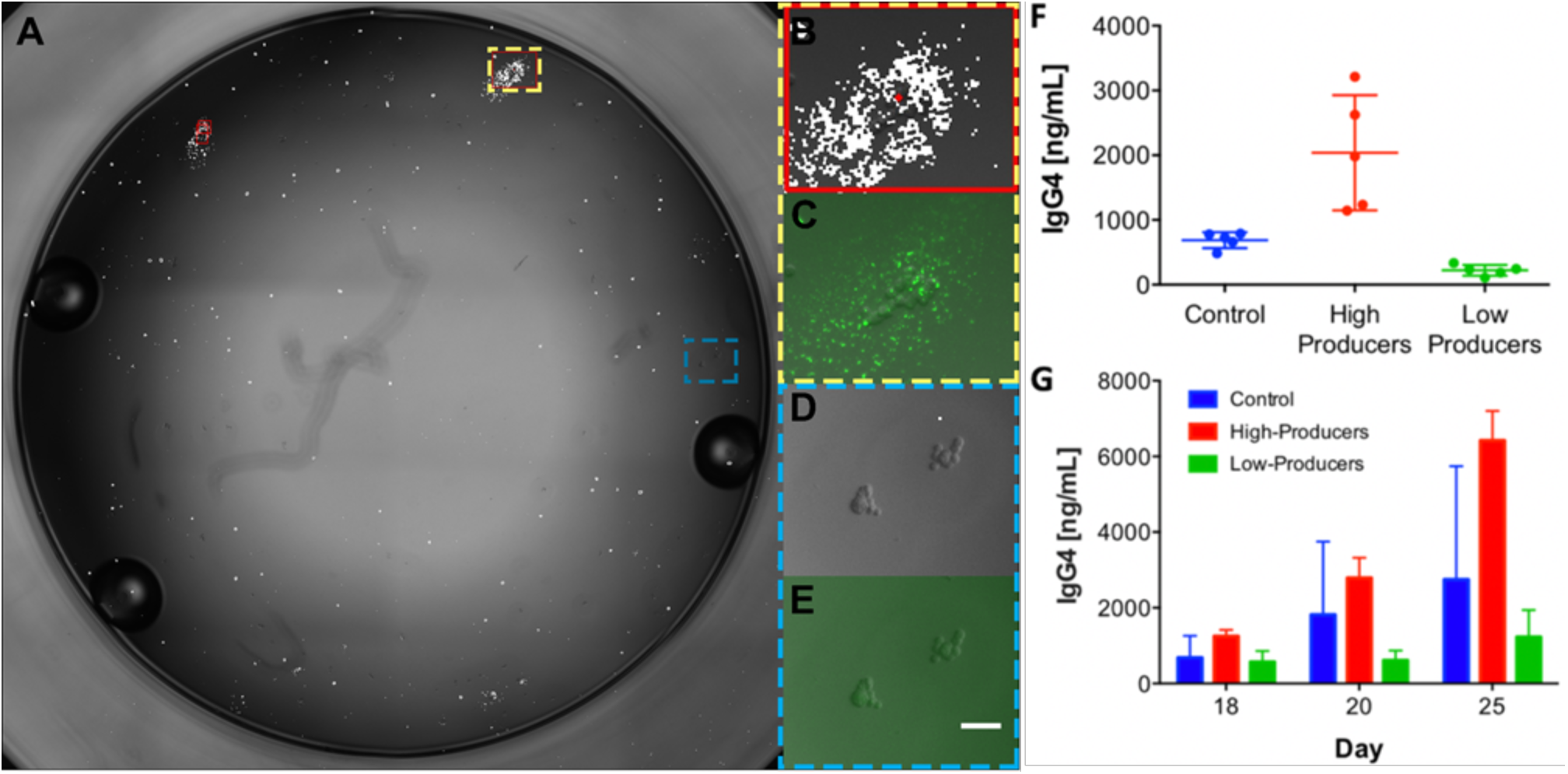
Immunofluorescence and ELISA-based assessment of antibody secretion. **A:** Brightfield image of a single well, seeded with antibody-secreting B13-24 CHO cells. Aggregates of secreted antibodies are annotated for visualization using white pixels. High-producing colonies are annotated using red rectangles. **B-E:** Enlarged annotated bright field (B,D) and fluorescence (C,E) images of high and low producing colonies. **F:** Secreted IgG4 concentration as measured by ELISA of supernatant sampled from the 96-well plate on day 15 (N = 5). **G:** High-producer cells maintained their increased IgG4 production during expansion up to 25 days into T-25 culture flasks, compared to control and low-producers. (Scale bar = 50 μm).

To maintain the selectively propagated cells, each well was supplemented with additional cell media every 2 days. The supernatant was sampled at regular intervals in order to measure the antibody concentration levels using ELISA (**Figure 3F-G**). The wells where the top three highest-producing colonies were selectively propagated exhibited a mean IgG4 concentration of 2039.3 ± 889.8 ng/ml (**Figure 4F**), which was a 2.96-fold increase in IgG4 production compared to control wells where no SLG was performed (N = 5, p = 0.0267). Correspondingly, wells where the three lowest-producing colonies were selectively propagated had a mean IgG4 concentration of 222.3 ± 83.4 ng/ml, which represented a 3.58-fold decrease in mean IgG4 concentration compared to control wells (N = 5, p = 0.0002). Together, these results demonstrated that SLG could be employed to selective propagate the high-secreting cells to increase the productivity of the culture.

We then sampled supernatant from the three groups of wells (high-producing, low-producing, and control) over 25 days of subsequent propagation in a subsequent experiment. As the cells grew to confluence, the culture was transferred from the 96-well plate into a 12-well plate, 6-well plate, and finally to a T-25 flask. Cell numbers in each well were counted on day 18 and normalized between each well prior to transfer to the 6-well plate and samples of the supernatant were obtained on days 18 (6-well plate), 20 (T-25 flask), and 25 (T-25 flask) for analysis using ELISA (**Figure 4G**). These results suggest that the high-producing cell line retained their high-level of IgG4 antibody production after 25 days of culture (N = 3, p < 0.1). The stability of the high-producing cell line is important in scale-up production since both volumetric and specific productivity declines as the cells age^39^. The large variation observed in the measured antibody secretion in the control well can be attributed to the inherent host cell heterogeneity, which can cause differences in growth rate and antibody productivity. Therefore, increasing the sample quantity will most likely not lead to reduced variability in antibody production. Significant differences between the average antibody production of the high-producers and controls are not anticipated, as the host cell line has undergone pre-selection such that the cells are capable of producing antibodies at high concentrations. However, it is expected that a small number of high producers, which surpass the production rate of the controls by a large margin, can be isolated.

## Discussion

In summary, our study leveraged the inverse thermal gelation property of methylcellulose to enable micro-patterning of hydrogels to selectively propagate cells based on features defined by imaging. We used this technology to screen antibody secretion from transgenic CHO cells. Using a fluorescence-labeled secondary antibody as an imaging probe, we developed custom software to measure and rank colonies based on their antibody secretion per cell. The top three highest-secreting colonies were allowed to propagate, while the remaining cells were encapsulated by hydrogel to arrest their growth. Hydrogel encapsulation via laser targeting prevents further cell growth because the phase transition from solution to gel hinders the cells from receiving oxygen and nutrients required for survival. This effect coupled with localized heating from the focused laser application leads to loss of viability of the target cells. The outgrowth of the selectively propagated cells showed a three-fold enhancement in antibody production over a non-selected sample. This production advantage was maintained when the outgrowth cells were expanded in culture for 25 days.

The ability to selectively propagate productive antibody-secreting cells could significantly reduce the timelines required for developing and testing antibody-based therapeutics. Typical screening by ELISA is done after 2-3 weeks of expansion from individual cell, when sufficient antibody has been secreted. Similarly, colony-picking from semisolid media requires 7-8 days of growth in order for colonies to grow to sufficient size for extraction by fine-needle aspiration^40^. As we showed here, the SLG process enables the antibody secretion rate to be measured and cells to be selectively propagated after only 1-2 days from plating.

A key advantage of the SLG process is the capability for secretion-based screening at the single-cell level in a standard growth environment. This capability is important for cell line development because clonally-selected cells exhibit significant phenotypic heterogeneity^41,42^. This phenotypic heterogeneity at the early stages of outgrowth could dramatically alter the productivity of the overall population. Specifically, low-producing cells use less energy for antibody expression and secretion, and are therefore able to devote more energy towards proliferation. The increased growth rate enables these cells to dominate the colony at the expense of high-producing cells^14,43^. When the producer cells are transferred to a bioreactor, these low-producing cells will outgrow the high-producing cells, resulting in an ultimate limit to productivity. The SLG process enables early stage screening at the single cell level in order to direct the culture towards high producers prior to initial colony expansion. It is possible to imagine devising algorithms to select for specific secretion and growth characteristics in multiple passes in order to direct the culture towards propagating more robust high-yielding cells.

The SLG process presents many characteristics that favor its integration into existing cell line development workflows. First, SLG uses standard reagents and consumables, thereby enabling users to preserve established workflows. Second, SLG is an image-based process that enables a single cell and its antibody secretion rate to be tracked as it develops into a colony. This capability can potentially be used to enable AI-powered screening to identify suitable antibody producing cells. Finally, screening using SLG is entirely non-contact. Fluid contact with the cell media occurs only when the cells are plated and when the cell population has been expanded after outgrowth. The non-contact property of the screening process prior to culture expansion of the outgrowth cells significantly reduces the potential for contamination and makes the process more amenable for automation.

Finally, it is possible to consider SLG as a general tool for image-based screening. In this context, specific phenotypes could be identified by imaging, and then SLG could be used to direct the growth of the culture towards a desired outcome. The ability to perform image-based cell selection enables rich phenotype screening that could potentially combine immunofluorescence and cytomorphological phenotyping in selective propagation of microbial cells, biologic producer cells, or even tissue cell subsets. Potential extension of this approach could also guide cell therapy, where undesirable phenotypes must be terminated prior to expansion and transplantation^44^.

## Materials and Methods

### Methylcellulose Media Preparation

To prepare a 2.5% methylcellulose solution, 950 g of distilled water and 24.81 g of Sigma methylcellulose powder (Cat # M0512, 4,000 cP, Sigma-Aldrich, St. Louis, MO) were autoclaved separately at 121°C for one hour. After autoclaving, using sterile technique, the methylcellulose powder was added to the distilled water while slowly stirring with a metal spoon. Magnetic stirring bars were added prior to sealing and the jar was placed on a magnetic stirrer and allowed to cool to room temperature (~20°C). After cooling, 17.7 g of Isocove’s Modified Dulbecco’s Medium (IMDM) powder (Sigma-Aldrich, St. Louis, MO) was added to the methylcellulose solution, stirred for an additional two hours and left overnight at 4°C.

To dilute to a final working concentration of 1% MC, 20 g of the 2.5% methylcellulose/IMDM solution, 23.7 mL of 1x IMDM media (ATCC, Manassas, VA), 5 mL heat-inactivated fetal bovine serum (FBS) (Thermo Fisher, Watham, MA), 800μL of 7.5% sodium bicarbonate solution (Thermo Fisher Scientific, Waltham, MA) and 500μL of 100X concentrated Penicillin/Streptomycin (P/S) solution (Thermo Fisher, Watham, MA) are mixed using a vortex mixer on high and left overnight at 4°C to disperse any bubbles. The next day, the methylcellulose/IMDM solution is filtered through a 0.45μm filter to remove any residual clumps and debris.

### Fluorescent cell lines

Fluorescent human bladder cancer cell lines, UM-UC13-EGFP and UM-UC3-mCherry, were provided by Dr. Peter Black at the Vancouver Prostate Centre at Vancouver General Hospital. These cells were maintained in Minimum Essential Medium (MEM) supplemented with FBS (10%v/v, Thermo Fisher, Watham, MA) and 1X P/S (v/v, Thermo Fisher, Watham, MA) in a humidified incubator at 37°C and 5% CO_2_. These cells were harvested using 0.25% Trypsin-EDTA (Thermo Fisher, Watham, MA) and centrifuged at 200 x g for 5 min. The harvested cells were mixed at a 1:1 ratio, resuspended in the same media supplemented with 1% methylcellulose, and plated in flat-bottom 96 well plates.

### Antibody-Secreting Cells

B13-24 (Cat # CRL-11397, ATCC, Manassas, VA) Chinese Hamster Ovary (CHO) cells were grown in IMDM (ATCC, Manassas, VA) containing 4 mM L-glutamine, 4500 mg/L glucose, and 1500 mg/L sodium bicarbonate supplemented with FBS (10%v/v) and 1X P/S (Thermo Fisher Scientific, Waltham, MA). B13-24 cells are capable of producing up to 100 μg/L of humanized immunoglobulin IgG4 at a rate of 25 pg/cell/day^38^.

### Plating in Methylcellulose Solution

To plate B13-24 cells in 1% methylcellulose /IMDM solution, 1000 cells were added to 1 mL of 1% MC/IMDM solution with 10 μL CloneDetect Human IgG (H+L) Specific, Fluorescein (Cat # K8202, Molecular Devices, Sunnyvale, CA). The solution was mixed slowly using a 1 mL syringe with 16g needle, taking care to prevent bubbles from forming and 100 μL of the solution was dispensed into the inner wells of a 96-well flat-bottom polystyrene plate (Cat# 353916, Corning Incorporated, Corning, NY). The 96-well plate was centrifuged at 530 x g for 5 minutes and incubated at 37°C, 5% CO_2_ for two days.

### Cell Culture and Supernatant Retrieval

Cells were incubated in a humidified incubator at 37°C, 5% CO_2_. After incubating in the methylcellulose/IMDM media for a varying amount of days based on the experiment, 200 μL of IMDM liquid media was added to each well containing cells. The supernatant was sampled and media refreshed at multiple time points to quantify the antibody level.

### Antibody Secretion Measurement by ELISA

Measurement of the antibody secretion produced by the grown cells was done using enzyme-linked immunosorbent assay (ELISA). An IgG4 Human ELISA Kit (Cat # BMS2095, Thermo Fisher, Waltham, MA) was purchased to verify the antibody secretion titer according to the manufacturers instructions.

### Statistical Analysis

Data was plotted in GraphPad Prism v7.0 (GraphPad Software, San Diego, CA) and analyzed using the two-tailed unpaired Student’s t-test with Welch’s correction. Data shown is Mean ± Standard Deviation.

### Image Analysis

Fluorescence microscopy images of UM-UC3 and UM-UC13 cells were analyzed using the image processing toolbox in Matlab 2018 to count the number of EGFP and mCherry pixels in the control and SLG-treated wells.

## Author Contributions

H.M. conceived the idea and supervised the work. J.C.Y.C., J.A.T., and K.M. performed the experimental work. J.H.L. contributed to the development of the optical system. All authors wrote the manuscript.

## Acknowledgments

We would like to acknowledge Peter Black for providing transgenic UM-UC13 and UM-UC3 cell lines used in this study. We would like to acknowledge Jeffrey Young, James Piret, and Manfred Koller for useful discussions related to this work. This work was supported by grants from Canadian Institutes of Health Research (381129, 426032), Natural Science and Engineering Research Council of Canada (2015-06541, 508392-17), and Michael Smith Foundation for Health Research (38714). H.M. acknowledges funding from the CIHR New Investigator Salary Award program (322375). J.H.L. acknowledges funding from the UBC Four Year Doctoral Fellowship program. J.A.T. and K.M. acknowledges funding from MITACS Accelerate program (IT13817, IT09621).

## References

1 J.-C. Lagier, S. Edouard, I. Pagnier, O. Mediannikov, M. Drancourt and D. Raoult, Clin. Microbiol. Rev., 2015, 28, 208–236.

2 S. K. Gupta and P. Shukla, Front. Pharmacol., 2017, 8, 419.

3 P. Gronemeyer, R. Ditz and J. Strube, Bioengineering, 2014, 1, 188–212.

4 C. M. Lopes-Ramos, J. N. Paulson, C.-Y. Chen, M. L. Kuijjer, M. Fagny, J. Platig, A. R. Sonawane, D. L. DeMeo, J. Quackenbush and K. Glass, BMC Genomics, 2017, 18, 723.

5 T. Lai, Y. Yang and S. K. Ng, Pharmaceuticals, 2013, 6, 579–603.

6 Z. Yang, S. Wang, A. Halim, M. A. Schulz, M. Frodin, S. H. Rahman, M. B. Vester-Christensen, C. Behrens, C. Kristensen, S. Y. Vakhrushev, E. P. Bennett, H. H. Wandall and H. Clausen, Nat. Biotechnol., 2015, 33, 842–844.

7 P. Ko, S. Misaghi, Z. Hu, D. Zhan, J. Tsukuda, M. Yim, M. Sanford, D. Shaw, M. Shiratori, B. Snedecor, M. Laird and A. Shen, Biotechnol. Prog., 2018, 34, 624–634.

8 S. M. Browne and M. Al-Rubeai, Trends in Biotechnology, 2007, 25, 425–432.

9 S. Dharshanan, H. Chong, C. S. Hung, Z. Zamrod and N. Kamal, Electronic Journal of Biotechnology,, DOI:10.2225/vol14-issue2-fulltext-7.

10 A. W. Caron, C. Nicolas, B. Gaillet, I. Ba, M. Pinard, A. Garnier, B. Massie and R. Gilbert, BMC Biotechnology, 2009, 9, 42–42.

11 J. M. Davis, J. E. Pennington, A. M. Kubler and J. F. Conscience, Journal of Immunological Methods, 1982, 50, 161–171.

12 S. M. Browne and M. Al-Rubeai, Trends in Biotechnology, 2007, 25, 425–432.

13 F. Gray, J. S. Kenney and J. F. Dunne, J. Immunol. Methods, 1995, 182, 155–163.

14 L. Hammill, J. Welles and G. R. Carson, Cytotechnology, 2000, 34, 27–37.

15 S. Carroll and M. Al-Rubeai, Expert Opinion on Biological Therapy, 2004, 4, 1821–1829.

16 E. G. Hanania, A. Fieck, J. Stevens, L. J. Bodzin, B. Palsson and M. R. Koller, Biotechnology and Bioengineering, 2005, 91, 872–876.

17 J. Kovac, Y. Gerardin and J. Voldman, Adv Healthc Mater, 2013, 2, 552–556.

18 T. Sun, J. Kovac and J. Voldman, Anal. Chem., 2014, 86, 977–981.

19 D.-S. Shin, J. You, A. Rahimian, T. Vu, C. Siltanen, A. Ehsanipour, G. Stybayeva, J. Sutcliffe and A. Revzin, Angew. Chem. Int. Ed. Engl., 2014, 53, 8221–8224.

20 K. J. Son, D.-S. Shin, T. Kwa, J. You, Y. Gao and A. Revzin, Lab Chip, 2015, 15, 637–641.

21 M. Shibuta, M. Tamura, K. Kanie, M. Yanagisawa, H. Matsui, T. Satoh, T. Takagi, T. Kanamori, S. Sugiura and R. Kato, J. Biosci. Bioeng., 2018, 126, 653–660.

22 M. Tamura, F. Yanagawa, S. Sugiura, T. Takagi, K. Sumaru, H. Matsui and T. Kanamori, Sci Rep, 2014, 4, 4793.

23 M.-P. Chien, C. A. Werley, S. L. Farhi and A. E. Cohen, Chem Sci, 2015, 6, 1701–1705.

24 L. Binan, J. Mazzaferri, K. Choquet, L.-E. Lorenzo, Y. C. Wang, E. B. Affar, Y. De Koninck, J. Ragoussis, C. L. Kleinman and S. Costantino, Nat Commun, 2016, 7, 11636.

25 B. Wognum and T. Lee, in Basic Cell Culture Protocols, eds. C. D. Helgason and C. L. Miller, Humana Press, Totowa, NJ, 2013, pp. 133–149.

26 T. G. Otsuji, J. Bin, A. Yoshimura, M. Tomura, D. Tateyama, I. Minami, Y. Yoshikawa, K. Aiba, J. E. Heuser, T. Nishino, K. Hasegawa and N. Nakatsuji, Stem Cell Reports, 2014, 2, 734–745.

27 A. W. Caron, C. Nicolas, B. Gaillet, I. Ba, M. Pinard, A. Garnier, B. Massie and R. Gilbert, BMC Biotechnology, 2009, 9, 42–42.

28 C. Lee, C. Ly, T. Sauerwald, T. Kelly and G. Moore, BioProcess International.

29 E. Heymann, Transactions of the Faraday Society, 1935, 31, 846.

30 J. Desbrières, M. Hirrien and S. B. Ross-Murphy, Polymer, 2000, 41, 2451–2461.

31 J. P. A. Fairclough, H. Yu, O. Kelly, A. J. Ryan, R. L. Sammler and M. Radler, Langmuir, 2012, 28, 10551–10557.

32 M. Takahashi, M. Shimazaki and J. Yamamoto, Journal of Polymer Science Part B: Polymer Physics, 2001, 39, 91–100.

33 A. W. Caron, C. Nicolas, B. Gaillet, I. Ba, M. Pinard, A. Garnier, B. Massie and R. Gilbert, BMC Biotechnology, 2009, 9, 42.

34 G. Bradski, Dr. Dobb’s Journal of Software Tools.

35 J. Canny, IEEE transactions on pattern analysis and machine intelligence, 1986, 8, 679–698.

36 N. Otsu, IEEE transactions on systems, man, and cybernetics, 1979, 9, 62–66.

37 S. Suzuki and K. A. be, Computer Vision, Graphics and Image Processing, 1985, 30, 32–46.

38 B13-24(ATCC CRL-11397), https://www.atcc.org/en/Products/Cells_and_Microorganisms/By_Tissue/Ovary/CRL-11397.aspx#generalinformation.

39 F. Li, N. Vijayasankaran, A. Shen, R. Kiss and A. Amanullah, mAbs, 2010, 2, 466–479.

40 S. Dharshanan, H. Chong, C. S. Hung, Z. Zamrod and N. Kamal, Electronic Journal of Biotechnology,, DOI:10.2225/vol14-issue2-fulltext-7.

41 S. L. Davies, C. S. Lovelady, R. K. Grainger, A. J. Racher, R. J. Young and D. C. James, Biotechnol. Bioeng., 2013, 110, 260–274.

42 L. M. Barnes, N. Moy and A. J. Dickson, Biotechnol. Bioeng., 2006, 94, 530–537.

43 Y. Kaneko, R. Sato and H. Aoyagi, J. Biosci. Bioeng., 2010, 109, 274–280.

44 C. A. Herberts, M. S. Kwa and H. P. Hermsen, J Transl Med, 2011, 9, 29.

